# Preprocessing Decisions Affect the Precision of P50 Estimates in the Paired-Click Paradigm

**DOI:** 10.64898/2026.07.22.739799

**Authors:** Evan Canny, Chris Paffen, Daan van Rooij, J. Leon Kenemans

## Abstract

Sensory gating is commonly indexed by suppression of the auditory P50 component in the paired-click paradigm. However, P50 findings have varied considerably across studies, laboratories, and populations, prompting calls for the standardisation of methodological practice. EEG preprocessing represents one source of methodological variation that requires standardisation. To date, filtering is the only preprocessing step to have been examined empirically. Segmentation and artifact handling have varied across paired-click studies without empirical evaluation, and, to our knowledge, the standardized measurement error (SME) has not been applied to P50 paired-click estimates. The present study addressed both gaps. Fifty-six neurotypical adults completed a paired-click paradigm consisting of 120 trials. Four preprocessing pipelines were constructed by crossing Segmentation (Short vs Long) with Artifact Handling (threshold-based rejection vs rejection of non-ocular artifacts followed by Gratton and Coles ocular correction). A bootstrapped SME (bSME) was used to quantify the precision of peak-to-peak P50 amplitudes at Cz and FCz, alongside an analysis of trial retention. Correction pipelines retained more trials and produced lower bSME values (higher precision) than Rejection pipelines, at both electrodes and across S1, S2, and suppression score outcomes. Segmentation and Artifact Handling did not operate independently: Segmentation had a negligible impact under Correction, whereas Long segmentation consistently produced the highest bSME values (lowest precision) under Rejection. A regression analysis showed that trial retention accounted for the large majority of the pipeline differences in bSME. This study therefore recommends that future paired-click studies correct, rather than reject, trials containing ocular artifacts, and adopt Short segmentation where matched S1–S2 averaging is not required.

**Impact statement:** Inconsistent P50 findings have long prompted calls for the standardisation of paired-click methodology, yet preprocessing decisions are rarely justified and, with the exception of filtering, have never been evaluated empirically. By quantifying the precision of P50 estimates, we show that artifact handling and segmentation affect data quality almost entirely through the trials they discard. Measurement precision, not convention or preference, should decide how P50 data are preprocessed.

## Introduction

Sensory gating refers to an internal filtering mechanism that enables the brain to regulate and prioritize incoming sensory information in order to prevent overload and preserve cognitive efficiency (Braff & Geyer, 1990). This mechanism functions as a form of pre-attentive inhibition, attenuating redundant or irrelevant inputs and reducing demands on attentional and processing resources (White & Yee, 2006; Venables & Maher, 1964; Micoulaud-Franchi et al., 2015, 2019). Sensory gating is commonly measured using auditory event-related potentials (ERPs), particularly the P50 component, in response to paired auditory stimuli (Adler et al., 1982).

In the paired-click paradigm, two identical auditory stimuli are presented in quick succession (typically 500ms apart), referred to as the first (S1) and second (S2) stimulus. The degree of P50 suppression is reflected by the reduction in P50 amplitude from S1 to S2, and is taken as an index of sensory gating effectiveness (Freedman et al., 1987; Clementz et al., 1998). Since Adler et al.’s (1982) seminal demonstration of reduced P50 suppression in schizophrenia, the paired-click paradigm has been widely used to characterize sensory gating across a range of neuropsychiatric and clinical populations, underscoring its value as an objective tool for studying early sensory processing.

However, despite its widespread use, paired-click P50 research findings have varied considerably across studies, laboratories, and populations, raising questions about the consistency and comparability of P50 suppression measures (de Wilde et al., 2007; Patterson et al., 2008). For example, studies show P50 suppression varies considerably within the healthy population, with reports showing P50 suppression can vary from strong levels (70-90% suppression) to “abnormal” levels (30-50% suppression) that are similar to schizophrenic patients (Kathmann & Engel, 1990; Lamberti et al., 1993; Patterson et al., 2008), a patient group for which P50 suppression deficits are considered a pathophysiological feature (Freedman et al., 2020). Findings from patient groups have also been inconsistent, such as for ADHD: some studies show P50 suppression deficits in ADHD patients compared to healthy controls (Holstein et al., 2013; Micoulaud-Franchi et al., 2015), while other studies have not replicated these findings (Lemvigh et al., 2020; le Sommer et al., 2020; Olincy et al., 2000).

Inconsistent research findings led to extensive investigation of P50 suppression metrics, where studies showed the psychometric reliability of P50 gating metrics range from poor to modest in neurotypical samples (Boutros et al., 1991; Clementz et al., 1997; Fuerst et al., 2007; Hall et al., 2006; Lamberti et al., 1993; Lu et al., 2007; Rentzsch et al., 2008; Smith et al., 1994) and schizophrenic patients (Clementz et al., 1997). Psychometric reliability refers to the proportion of total between-participant variance in a P50 score that is attributable to true individual differences rather than measurement error (Clayson et al., 2021; Luck et al., 2021, Supplement S1). In P50 paired-click studies, psychometric reliability is assessed using the intraclass correlation coefficient (ICC) and test-retest correlations. Researchers have explored the possibility that methodological heterogeneity may contribute to variation in findings and P50 measures across studies (de Wilde et al., 2007; Patterson et al., 2008).

Several methodological factors have varied considerably across paired-click studies that may have influenced the assessment of P50 suppression. Of these factors, EEG preprocessing decisions represent one class of methodological variation that may plausibly contribute to differences in P50 findings. Among these preprocessing decisions, filtering is the only step that has received direct empirical examination (Jerger et al., 1992). de Wilde et al.’s (2007) review of the schizophrenic P50 literature showed effect sizes were associated with high-and low-pass filter settings, while Patterson et al., (2008) showed that high-pass filter setting (0.8-vs 10-vs 30 Hz-high-pass) significantly affected P50 ratios across four healthy control groups. Extending this, Chang et al. (2012) demonstrated that filter settings can directly influence P50 amplitude and ratios, signal-to-noise ratio, and variability, with certain bandpass ranges (e.g., 10 Hz high pass filter) associated with more stable ratio estimates and higher signal-to-noise ratio (SNR). These findings suggest that preprocessing decisions can affect P50 measures. However, it remains unclear whether preprocessing decisions other than filter settings also impact P50 measures. The present study focuses on two preprocessing steps that have varied across paired-click studies and have yet to be empirically examined: segmentation and artifact handling.

Segmentation and artifact handling procedures are implemented prior to averaging and therefore directly influence how and which trials are retained for analysis. As a result, differences in segmentation and artifact handling decisions may affect the number of trials contributing to the averaged P50. This may be important because trial count is positively related to the SNR of event-related potentials (ERPs), such that increasing the number of trials improves the SNR of the averaged signal (i.e., SNR increases as a function of the square root of the number of trials; Luck, 2014). Beyond its effects on signal quality, trial count also plays a critical role in determining the statistical power (the probability of detecting a true effect) of ERP analyses. That is, increasing the number of trials increases the likelihood of detecting true effects, particularly when effect sizes are small or noise levels are high (Boudewyn et al., 2018). Therefore, methodological decisions that influence the number of trials that are retained for averaging, especially segmentation and artifact handling, should be chosen carefully on the grounds of empirical support. This logic is particularly pertinent when analysing low-amplitude and short-latency ERPs, such as the P50 component (∼1–3 µV; ∼50ms onset time), as the signal of interest is more easily influenced by voltage fluctuations that are not time-locked to the stimulus.

Furthermore, and more specifically relating to the P50 ERP component, multiple studies have previously demonstrated that increasing the trial count is associated with improvements in the psychometric reliability of P50 measures. Boutros et al. (1991) showed that doubling trial numbers increased test–retest reliability of P50 ratio scores. These findings were replicated and extended by controlling within-session, time-related effects (Clementz et al., 1997; Dalecki et al., 2011). Collectively, this work suggests that reliable estimation of P50 measures requires relatively large numbers of trials, with recommendations typically ranging from 40 trials as a minimum threshold to at least 100 trials to maximise within-subject stability (Boutros et al., 2004, 2009; Dalecki et al., 2011; Fuerst et al., 2007).

In sum, segmentation and artifact handling represent two preprocessing steps that may plausibly influence P50 measures. This methodological focus aligns with recent ERP research that has emphasized the use of objective, quantitative approaches to evaluate and guide EEG processing decisions and their impact on ERP data quality (Zhang et al., 2024a, 2024b, 2024c; Zhang & Luck, 2023). However, before discussing the ERP data-quality approach led by Luck and colleagues, it is first necessary to establish how the approaches to artifact handling and segmentation preprocessing steps have been implemented across the paired-click literature so far.

### Segmentation

Paired-click studies have adopted two approaches to EEG segmentation, which can be referred to as single and dual segmentation. In single segmentation^1^, separate short epochs (-100 to 200/400ms) are created relative to the S1 and S2 markers, yielding two independent segments per trial. In contrast, dual segmentation^2^ involves extracting a single, longer epoch (-200ms to 1000/1200ms) time-locked to S1 that contains both the S1 and S2 responses within the same segment. Because segmentation precedes artifact rejection, the choice of segmentation strategy can have direct consequences on the number of trials contributing to the averaged ERP.

In dual segmentation, an artifact occurring anywhere within the longer epoch, leads to rejection of the entire paired-click trial, resulting in matched S1–S2 averages but reduced trial counts. In contrast, under single segmentation, S1 and S2 segments are rejected independently, which can increase trial retention but may result in unmatched S1 and S2 averages. One way to avoid this mismatch is to reject paired trials jointly, such that rejection of either the S1 or S2 segment leads to rejection of the corresponding segment from the same trial. Although this approach has been used (e.g., Lijffijt et al., 2009), it is not common practice.

The lack of consistency in segmentation indicates that no standard method has been adopted across the paired-click literature, and the consequences of this variability for trial retention remain unclear. Given the established relationship between trial retention, signal-to-noise ratio, and the psychometric reliability of P50 measures (Boutros et al., 1991; Clementz et al., 1997; Dalecki et al., 2011), differences in segmentation strategy may plausibly influence downstream P50 measures, and therefore warrant closer examination.

### Artifact handling

Artifact handling procedures have also differed across paired-click studies. The most common approach is threshold-based artifact rejection, where trials exceeding a fixed amplitude criterion (±75 µV or ±100 µV) at EEG and EOG channels are excluded. This procedure removes trials containing eye blinks or eye movements prior to averaging, which can reduce trial retention and, in some cases, lead to exclusion of participant averages that fail to meet minimum trial requirements (Fuerst et al., 2007). A second approach applies artifact rejection to EEG channels only to remove non-ocular artifacts, and then applies ocular correction methods on the remaining trials to correct for ocular artifacts (eye blinks). Because eye blinks are corrected rather than rejected, this combined rejection-correction approach retains more trials for averaging than threshold-based approaches, and is therefore likely to produce systematic differences in trial counts between the two artifact handling procedures.

Although paired-click studies vary in their approach to artifact handling, few studies report the number of trials removed during preprocessing or provide explicit justification for the choice of artifact handling strategy (for a notable exception, see Hall et al., 2006).

Furthermore, researchers have previously recommended against the use of threshold-based artifact rejection as a means to handle ocular artifacts (Croft & Barry, 2000). Therefore, given the established relationship between trial retention, signal-to-noise ratio, and the psychometric reliability of P50 measures, differences in artifact handling may plausibly influence downstream P50 measures, and, next to segmentation strategy, warrant closer examination.

### Data quality metric – Standardized measurement error

Recent ERP methodology work has increasingly emphasized the use of objective, quantitative metrics to evaluate the impact of preprocessing decisions on data quality (Zhang et al., 2024a, 2024b, 2024c; Zhang & Luck, 2023). One such metric is the standardized measurement error (SME), introduced by Luck et al. (2021) as an index of data quality for averaged ERP components. The SME quantifies the precision of an ERP amplitude or latency score. Precision refers to the degree to which repeated measurements of the same participant under unchanged conditions would yield similar results (Luck et al., 2021). Currently, the paired-click literature has assessed P50 data quality using indices of psychometric reliability. However, psychometric reliability is driven by both between-participant true-score variance and within-participant measurement error, and therefore reflects both sample composition and per-participant data quality (Luck et al., 2021, Supplement S1). Precision, in contrast, is a property of an individual participant’s averaged ERP score and is independent of between-participant variability (Clayson et al., 2021). Larger SME values indicate lower precision, reflecting greater noise-related variability in the score. Importantly, the SME is sensitive to factors that produce imprecision in ERP scores, such as the amount of noise in the single-trial EEG and the number of trials contributing to the averaged waveform (Luck et al., 2021). As outlined above, segmentation strategy and artifact handling both influence the number of trials retained for averaging. These differences in preprocessing pipelines are therefore expected to directly affect the SME.

Previous evaluations of P50 measures in the paired-click literature have used the intraclass correlation coefficient (ICC; e.g., Rentzsch et al., 2008) and a within-subject standard deviation across block averages (WSD; e.g., Dalecki et al., 2011). As Luck et al. (2021) point out, a suitable data quality metric for averaged ERPs must satisfy three criteria: (a) it must be computable independently for each participant; (b) it must quantify the extent to which noise impacts the actual amplitude score used in the statistical analysis; and (c) it must represent the precision of that score.

The ICC is a group-level index of psychometric reliability that produces a single estimate for the entire sample, failing criterion (a). Its value is also determined by between-participant true-score variance and within-participant measurement error, so the ICC does not isolate the noise impact on participant-level scores, failing criterion (b) (Luck et al., 2021, see supplement S1). The WSD is computed by scoring the P50 peak amplitude from each block average separately and taking the standard deviation across those block-level scores (Dalecki et al., 2011). Because block averages contain substantially more residual noise than the full session average, the peak amplitude scored from each block is noise-distorted. As a result, the WSD does not estimate the precision of the peak amplitude derived from the full averaged waveform, failing criterion (b). The SME overcomes these limitations, making it the more suitable index of averaged ERP data quality.

The SME has two variants: the analytic SME (aSME) and the bootstrapped SME (bSME). The aSME applies the standard error formula to single-trial mean amplitude scores within a specified time window, and is suitable for time-window mean amplitude scoring (Luck et al., 2021). For peak-based scores such as the P50, the aSME shares the same structural flaw as the WSD. Peak amplitudes scored from single trials are noise-distorted relative to the peak scored from the full averaged waveform. Therefore, the standard deviation across single-trial peak scores does not estimate the precision of the peak that enters the statistical analysis, failing criterion (b). The bSME, described next, avoids this flaw by using bootstrapped averages of the full trial set.

The bSME uses a bootstrapping procedure, in which trials are repeatedly resampled with replacement from the full set of trials to simulate new averaged waveforms. On each of thousands of iterations, a new averaged waveform is constructed from the resampled trials, from which the peak amplitude is then scored. Because each resampled average is based on the full trial count, it contains substantially less noise than a single-trial or block-level average. The peak scores derived across iterations are therefore directly comparable to the peak scored from the actual averaged waveform. The standard deviation of those bootstrap peak scores validly estimate the precision of the score that enters the statistical analysis. The bSME is therefore computed independently for each participant (satisfying criterion a), quantifies the noise impacting the peak amplitude score used in the statistical analysis (satisfying criterion b), and represents the precision of that score (satisfying criterion c). It is therefore the appropriate metric for evaluating whether preprocessing pipeline decisions improve or reduce the data quality of averaged P50 peak amplitudes (Luck et al., 2021).

Recent studies have demonstrated this utility empirically, showing that preprocessing choices can influence SME values across a range of ERP components (Zhang et al., 2024a; Zhang et al., 2024b; Zhang et al., 2024c; Zhang & Luck, 2023). Specifically, Zhang and Luck (2023) used the SME to quantify the data quality of 7 commonly studied ERP components across a number of paradigms and scoring methods, providing a benchmark of data quality for other laboratories. Zhang et al. (2024a; 2024b; 2024c) extended this framework by using the SME to evaluate the impact of filter settings and artifact rejection methods on the data quality of those same 7 ERP components, demonstrating that both preprocessing decisions influence ERP amplitude score quality.

To date, the SME has not been applied to the P50 paired-click paradigm, and the metrics previously used to evaluate P50 measures in this literature, the ICC and WSD, do not meet the criteria for a data quality metric of averaged ERPs. In response to calls for greater standardisation of methodological practices in the paired-click literature (de Wilde et al., 2007; Patterson et al., 2008), the present study addresses both gaps by applying the bSME to empirically examine two preprocessing choices that have varied considerably across studies: segmentation strategy and artifact handling. The aim is to assess their impact on trial retention and P50 data quality.

## Methods

### Sample

56 individuals participated in this study (female N = 48, age M = 24.79 years, SD = 5.59 years). All participants were aged between 18 and 44 years, had no psychiatric, neurological, or hearing conditions, and had no prior exposure to the stimuli. Due to the known effects that substances have on neural and attentional processes (Böcker et al., 2010; Dockree et al., 2017; Gilbert et al., 2000), participants were instructed to abstain from smoking, caffeine, alcohol, stimulants, or other drugs after 22:00 on the night before participation. All participants provided written informed consent prior to participation, and the study was approved (25-0352) by the Ethics Committee of the Faculty of Social and Behavioural Sciences, Utrecht University.

### Paradigm

The paired-click paradigm was presented using *PsychoPy* (Peirce et al., 2019) with auditory stimuli delivered through *PsychPortAudio*. Each trial consisted of a pair of auditory clicks (S1 and S2), each 20ms in duration (1000 Hz pure tone; 4ms rise/fall time), separated by a 500ms interstimulus interval (ISI) and followed by a variable 3–4 s interpair interval (IPI; Rentzsch et al., 2008). A total of 120 paired-click trials were presented across three blocks of 40 trials each. The tones were presented binaurally at 100 dB SPL (see Appendix E for description of intensity calibration procedure) through ER-3C insert earphones (Micoulaud-Franchi et al., 2015), a level sufficient to elicit robust P50 components (De Wilde et al., 2007; Holstein, 2011). The intensity at the level of the ear canal was likely closer to 80 dB, as the same stimulus read 80 dBA when the airtight seal was removed during calibration. Before the task began, participants were familiarized with the tones to ensure they did not evoke a startle response. During EEG recording, participants watched a silent movie presented (EIZO FlexScan S1932 monitor) at the centre of the screen. Participants were instructed to focus on the movie throughout the task to control for attentional allocation across individuals (Dalecki et al., 2015, 2016). After each block, participants answered a short multiple-choice question about the movie and feedback was provided. If participants answered the question incorrectly, they were instructed to pay closer attention to the movie. The paired-click task was administered as part of a broader experimental protocol containing two additional tasks, which will be reported elsewhere. The order of the tasks was counterbalanced across participants to minimize potential confounding effects of arousal (White & Yee, 2006).

### EEG recording

The EEG session took place in a dimly lit, sound-attenuated room. Participants were seated comfortably in front of the stimulus display and used a chin rest to minimize head movements throughout the recording.

EEG data were recorded using a 64-channel *BioSemi ActiveTwo* system (BioSemi B.V., Amsterdam), with the Common Mode Sense (CMS) and Driven Right Leg (DRL) electrodes serving as the ground and reference. Electrodes were placed according to the 10– 10 system, including Fpz, AFz, Fz, FCz, Cz, CPz, Pz, POz, Oz, FP1/2, AF3/4/7/8, F1/2/3/4/5/6/7/8, FC1/2/3/4/5/6, FT7/8, C1/2/3/4/5/6, T7/8, CP1/2/3/4/5/6, TP7/8, P1/2/3/4/5/6/7/8, PO3/4/5/6/7/8, and O1/2. Vertical electrooculogram (EOG) activity was recorded using two electrodes placed ∼1cm above and below the left eye, and horizontal EOG was recorded using electrodes placed ∼1cm beyond the outer canthi. Two electrodes were placed on the mastoids for offline re-referencing. Data were sampled at 2048 Hz and recorded using BioSemi ActiView (version 9.00) software with an online low-pass filter of 417 Hz.

### Preprocessing pipelines

Data processing was carried out on BrainVision Analyzer (Version 2.3, Brain Products GmbH, Gilching, Germany). Several preprocessing steps were applied uniformly across all pipelines: EEG data were downsampled to 500 Hz (after applying a 225 Hz, 24 dB/octave anti-aliasing filter), re-referenced to the average reference^3^, and 10-50-Hz bandpass filtered using zero-phase Butterworth filters (12dB/octave), along with a 50Hz notch filter. All data were baseline corrected to the prestimulus period (-100 to 0ms relative to S1 or S2) before artifact handling^4^. These are all standard preprocessing steps applied throughout the P50 paired-click literature.

The four preprocessing pipelines differed along two dimensions: the segmentation strategy (single or dual segmentation) and the artifact handling method (threshold-based or artifact rejection/correction). Single segmentation involved creating separate short epochs around the S1 and S2 markers (for P50:-100 to 300ms). Dual segmentation involved creating a longer epoch relative to the S1 marker so that both S1 and S2 responses were contained within the same segment (-100 to 1000ms). The dual segmentation approach entails an additional 300ms that may contain non-EOG artifacts and result in the loss of more trials compared to single segmentation. This could lead to improved SNR using the single segmentation approach.

Artifact handling was carried out either using a fixed ±75μV threshold applied to all EEG and EOG channels or using an artifact rejection/correction procedure, in which non-ocular artifacts were removed at midline electrodes (Fz, FCz, Cz, CPz, Pz) using gradient, amplitude, and low-activity criteria^5^, followed by Gratton and Coles’ ocular correction method to correct for ocular artifacts. This yielded four preprocessing pipelines:

These pipelines represent common combinations of segmentation and artifact handling strategies used in paired-click studies and provide a structured basis for evaluating their impact on the quality and interpretability of the resulting S1 and S2 averages. All P50 analyses were conducted at Cz and repeated at FCz. Cz was selected because it is the electrode most commonly used for P50 quantification in the paired-click literature (Clementz et al., 1998), whereas FCz was additionally examined because the S1 P50 component reached its largest amplitude at this electrode in the present dataset. Figure 1 (below) represents the grand average localizer (collapsed across S1 and S2) showing the P50 component at Cz (left) and FCz (right) for each preprocessing pipeline.

**Figure 1.**
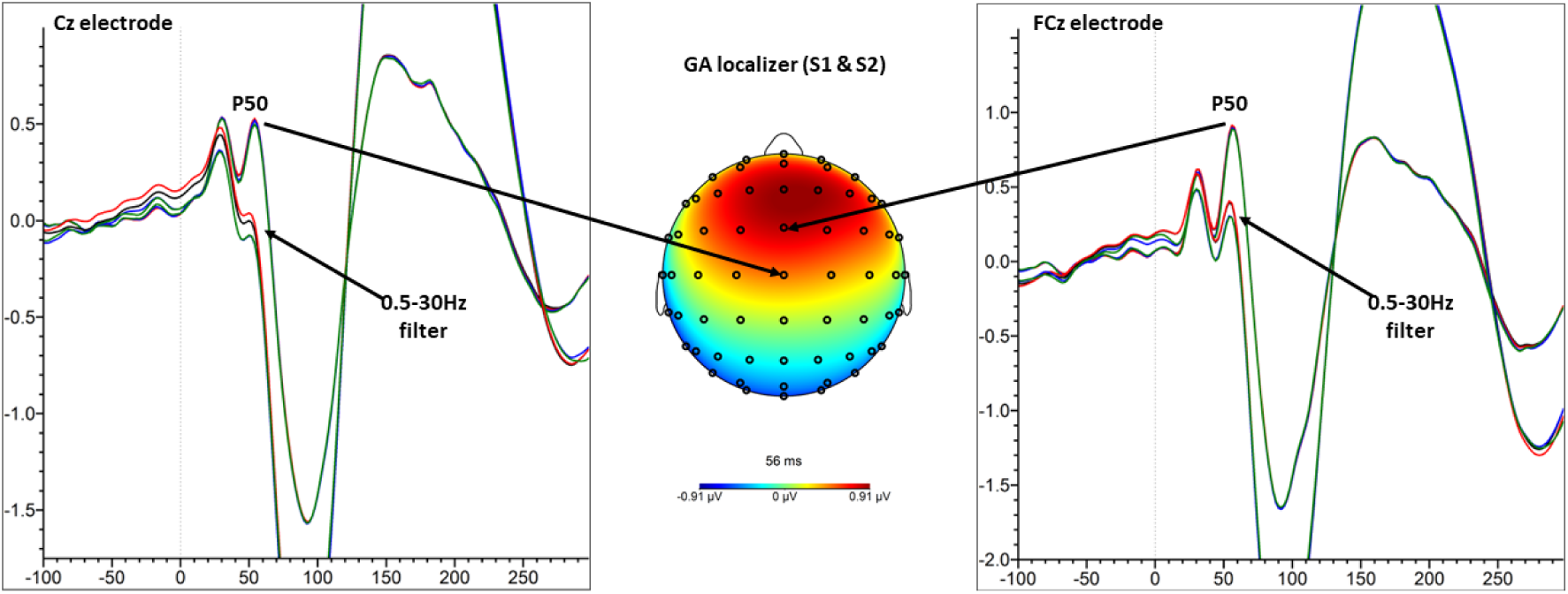
Grand Average Localizer (S1 & S2) P50 at Cz and FCz Electrodes. *Note.* This figure demonstrates the grand average (GA) localizer (S1 & S2 averages) at Cz and FCz, along with the topographical distribution at the P50 peak (56ms). 0ms is click onset. All four preprocessing pipelines are shown: Short Correction (black), Long Correction (red), Short Rejection (blue), Long Rejection (green). Given the high similarity of the 4 pipelines at the grand average level, the topographical distribution map is taken from the Short Correction pipeline only. The waveforms are also shown when a 0.5-30Hz filter is applied, significantly reducing the P50 component and increasing the N100 and P200 components.

For all analyses, P50 amplitudes (S1 P50 and S2 P50) and derived gating metrics (difference scores: S1 – S2; ratio scores: S2/S1) were quantified using peak-to-peak (P50 – N40) rather than the prestimulus baseline-to-peak assessment. Peak-to-peak will be abbreviated as “pp” (e.g., S1 P50pp). This is the most commonly used approach in paired-click studies for several methodological reasons. First, the N40 has been shown not to exhibit sensory gating effects, making it a suitable reference for estimating the magnitude of gated components (Arnfred et al., 2001). Second, the prestimulus period can contain lingering neural activity from the previous stimulus, such as contingent negative variation, that can interfere with the prestimulus period (Smith et al., 1994). Third, the P50 can be affected by the subsequent, much-larger, N100 component, which can drag down the P50 relative to the prestimulus period (Jerger et al., 1992). Although P50-specific bandpass filtering settings have been established to reduce this constraint (Chang et al., 2012), peak-to-peak amplitude assessment remains the most common approach utilised by P50 paired-click studies.

In the present dataset, inspection of the grand average waveforms provided an additional, empirically grounded reason to adopt the peak-to-peak approach. The grand average waveform (see figure 2 below) shows that the S1 N40 and S2 N40 show a similar latency profile (peak at ∼42ms). However, there is a clear amplitude difference when the waveforms are baseline corrected to the prestimulus period (-100 to 0ms relative to S1 or S2). As such, baseline-to-peak assessment would not provide a fair or valid comparison between S1 P50 and S2 P50 amplitudes, and would artificially inflate the derived gating metrics. Therefore, peak-to-peak assessment of P50 amplitudes provides a more valid analysis of P50 amplitudes and the derived gating metrics.

**Figure 2.**
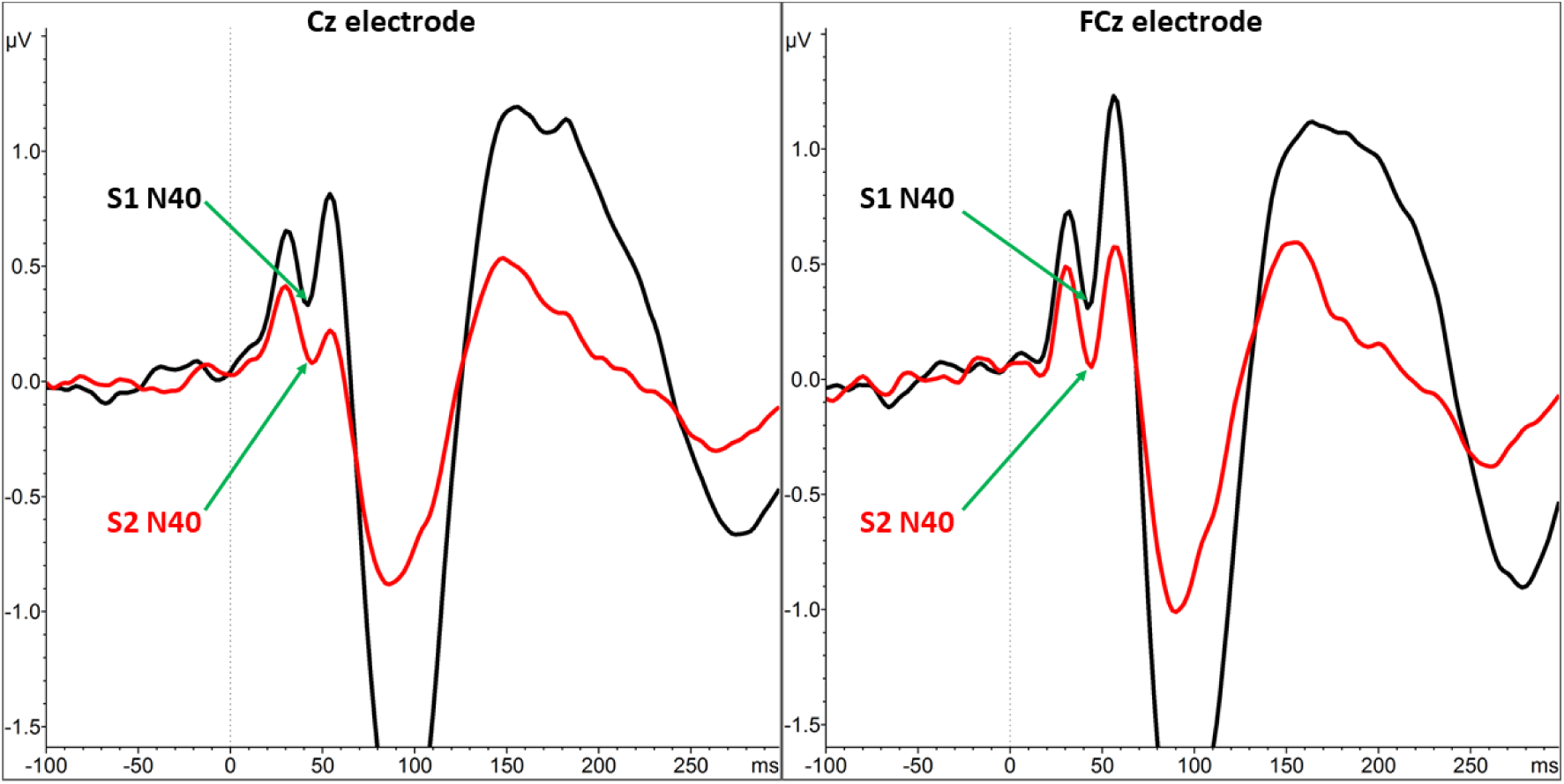
Grand Average S1 and S2 Waveforms at Cz and FCz Electrodes. *Note.* This figure demonstrates the S1 (black) and S2 (red) grand average (GA) collapsed across all 4 preprocessing pipelines (i.e., 4 pipelines collapsed into a single grand average for S1 and S2 separately) at Cz (left) and FCz (right). The figure shows the difference between the S1 N40 and S2 N40 at both electrodes. Green arrows are used to direct the reader to the S1 N40 and S2 N40. 0ms is click onset.

**Figure 3.**
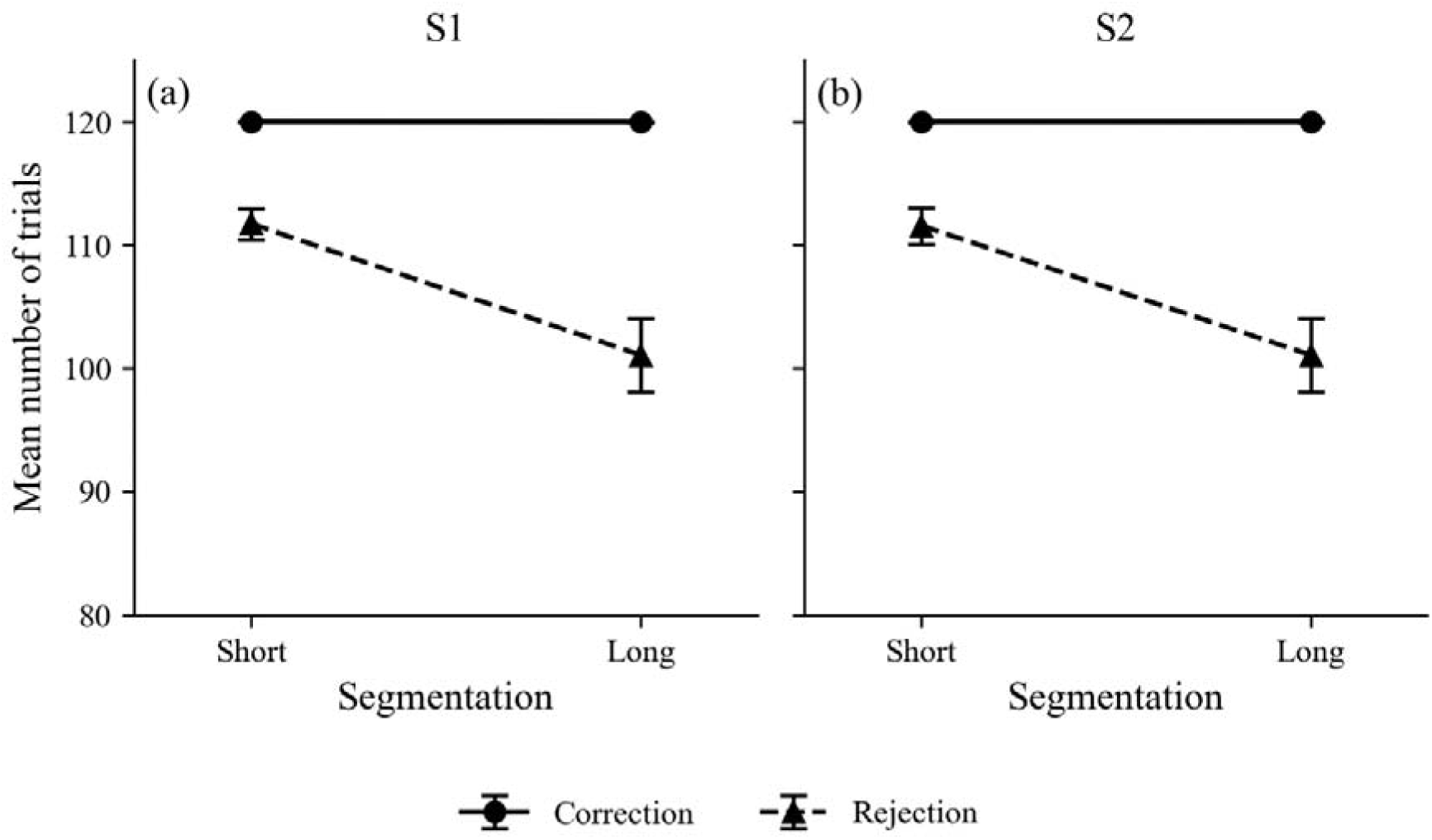
Mean Number of Trials Retained for S1 and S2 Across Preprocessing Pipelines. *Note.* Mean trial counts for S1 (panel a) and S2 (panel b) across Segmentation levels (Short vs Long) separately for Correction and Rejection artifact handling strategies. Error bars represent the standard error of the mean trial count within each pipeline.

**Figure 4.**
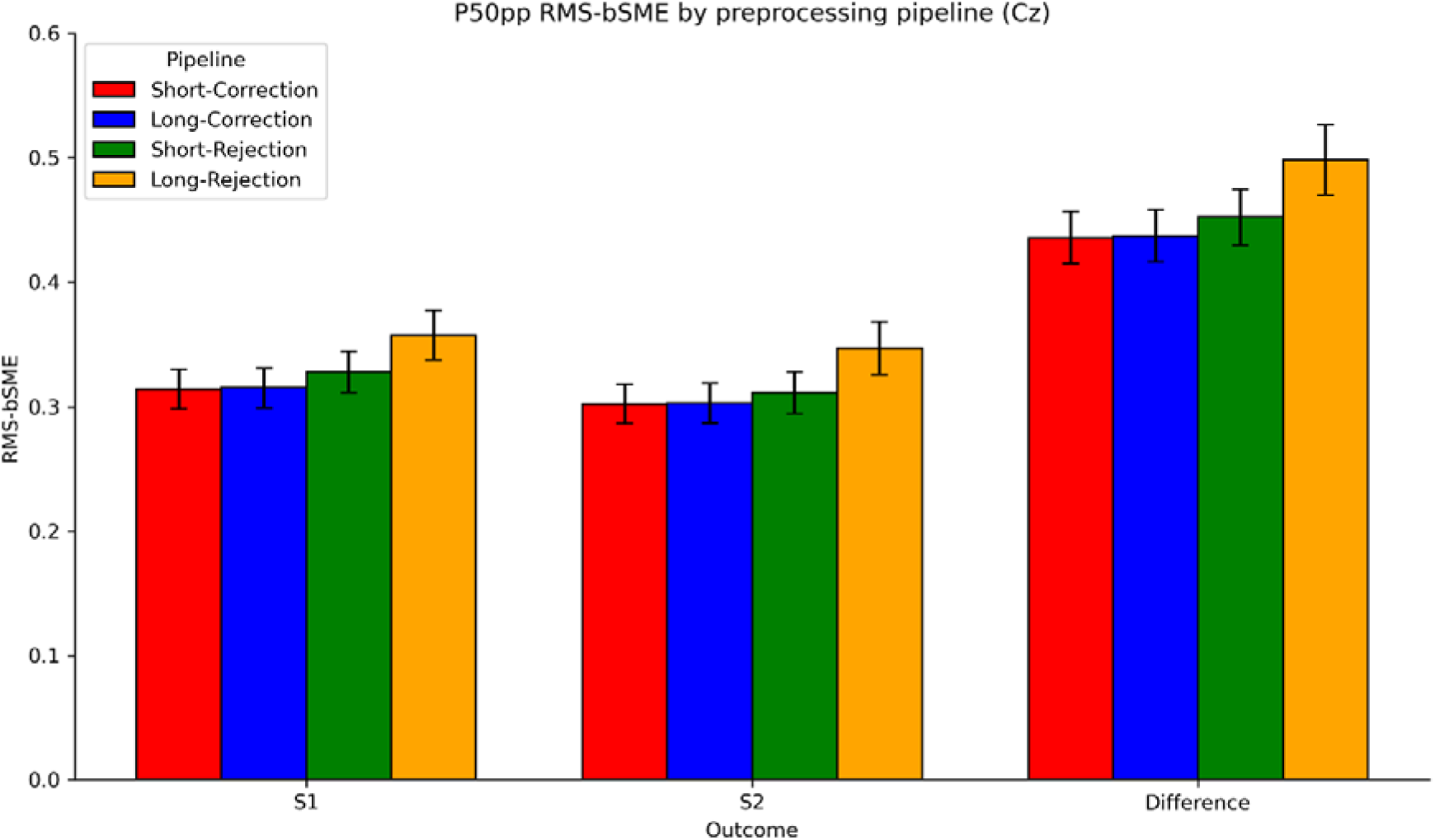
RMS-bSME P50pp values at Cz across all pipelines. *Note.* All pipelines are shown: Short Correction (red), Long Correction (blue), Short Rejection (green), and Long Rejection (orange). Error bars represent the standard error of the RMS-bSME for each pipeline.

### Analyses Overview

To evaluate the effects of preprocessing pipeline on the quality and reliability of paired-click P50 data, analyses were conducted on trial retention and standardized measurement error (SME). First, trial retention was examined to characterise how the four preprocessing pipelines differ in the number of trials retained for averaging. Second, the bSME was used to assess how the pipelines differ in P50 data quality, with lower bSME values indicating greater measurement precision. Third, a regression analysis was conducted to examine the extent to which any bSME differences between pipelines are explained by differences in trial retention, given that the SME is sensitive to the number of trials contributing to the averaged waveform (Luck et al., 2021).

### Trial retention

To examine whether trial retention differed across preprocessing pipelines, the four pipelines were treated as the four cells of a 2 × 2 within-subject factorial design, with Segmentation (Short vs Long) and Artifact Handling (Correction vs Rejection) as the two factors. The main effects and interaction were tested using three planned orthogonal contrasts. For each participant, contrast scores were computed by applying contrast weight vectors to their raw trial counts across the four pipelines: an Artifact Handling main effect contrast (Correction minus Rejection, averaged across segmentation levels), a Segmentation main effect contrast (Short minus Long, averaged across artifact handling levels), and a Segmentation × Artifact Handling interaction contrast. Each contrast score was tested against zero across participants.

Prior to inferential testing, the normality of each contrast score distribution was evaluated using Shapiro-Wilk tests. Where normality was met, a one-sample t-test was used; where normality was violated, a one-sample Wilcoxon signed-rank test was used instead, with rank-biserial correlation reported as the effect size measure (*r*_rb; Kerby, 2014). Confidence intervals for the rank-biserial correlation were obtained using a percentile bootstrap, in which participants were resampled with replacement 10,000 times and the effect size was recomputed on each resample, with the 2.5th and 97.5th percentiles of the resulting distribution taken as the lower and upper bounds. Holm correction was applied across the three contrast p-values within each epoch, as pairwise Spearman correlations between the three contrast score distributions confirmed they were not independent.

### Standardized measurement error

The following sections provide a detailed description of the computational procedures used to derive SME estimates, in order to ensure transparency and reproducibility of the analysis. A bootstrap-based SME (bSME) was used to assess the precision of peak-to-peak P50 amplitudes (Luck et al., 2021).

### bSME

For each participant, preprocessing pipeline, and stimulus (S1 and S2), single-trial segments were resampled with replacement and averaged to form bootstrap ERPs. This procedure was repeated 10,000 times per condition. From each accepted bootstrap ERP pair, peak-to-peak P50 amplitudes were derived at Cz and FCz using a peak-picking algorithm (see following paragraph for description of algorithm). Peak-to-peak P50 amplitude was defined as the difference between the P50 peak and the preceding N40 trough (P50 − N40).

Peak selection was performed using fixed, physiologically plausible latency constraints. The P50 peak was identified first as the most positive local extremum within a latency window of 40–80ms post-stimulus. Local extrema were defined using a neighbourhood rule (k = 3 samples on each side): candidate peaks were required to exceed the mean of their neighbouring samples. If no local extremum satisfied this criterion, the maximum value within the window was selected as a fallback. The N40 trough was then identified as the most negative local extremum within a window spanning 25ms up to, but excluding, the selected P50 latency, using an analogous neighbourhood rule in which candidate troughs were required to be lower than the mean of the neighbouring samples, with a minimum-value fallback if needed. Bootstrap iterations were accepted on the basis of structural validity only, such that a valid P50 had to be identified within the canonical P50 window and a valid N40 had to occur before the selected P50 latency.

To preserve the paired-click structure, S1 and S2 bootstrap ERPs were generated and evaluated jointly: a bootstrap iteration was accepted only if structurally valid component estimates were obtained for both S1 and S2; otherwise, the iteration was discarded and resampled, with up to 50 attempts permitted per bootstrap slot. For each accepted bootstrap iteration, peak-to-peak amplitudes were first computed separately for S1 and S2 (S1 P50pp, S2 P50pp). Peak-to-peak amplitudes below 0.01 µV were set to 0.01 µV. Suppression indices were derived for the difference score (S1 − S2) and ratio score (S2/S1).

For each participant, preprocessing pipeline, and outcome, the bSME was computed as the standard deviation of the scores obtained across bootstrap iterations:

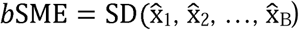

Where 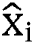 denotes the peak-to-peak amplitude score derived from the *i*th bootstrap averaged ERP waveform and *B* = 10,000 denotes the number of bootstrap iterations. Analyses were conducted separately for S1 and S2 peak-to-peak amplitudes, as well as for derived measures including the S1–S2 difference score and ratio score.

#### RMS-bSME: Group-Level Data Quality

To summarise data quality at the group level, the root mean square SME (RMS-SME) was computed for each preprocessing pipeline and outcome of interest by squaring each participant’s bSME value, averaging the squared values across participants, and taking the square root. This aggregation method is preferred over the arithmetic mean because RMS-SME is more directly related to effect size and statistical power (Luck et al., 2021; Zhang et al., 2024a). In line with Zhang et al., (2024a), a bootstrap standard error was computed for each RMS-SME estimate using 10,000 resamples of the participant-level bSME values. RMS-SME values and their bootstrap standard errors are presented separately for Cz and FCz as grouped bar charts, with S1, S2, and the difference score as three outcome groups on the x-axis and four adjacent bars within each group representing the four preprocessing pipelines.

#### Factorial Decomposition of Pipeline Differences

To decompose the observed pipeline differences into their factorial components, the outcomes of the four preprocessing pipelines were treated as the four cells of a 2 × 2 within-subject factorial design, with Segmentation (Short vs Long) and Artifact Handling (Correction vs Rejection) as the two factors. For each participant and outcome, the four pipeline bSME values were first squared to move them from the SD-like scale of the raw SME onto a variance-like scale.

Three planned orthogonal contrast scores were then computed on the squared values using fixed weight vectors applied in the pipeline order Short-Correction, Long-Correction, Short-Rejection, Long-Rejection: an Artifact Handling main effect contrast with weights [1, 1, −1, −1], representing Correction minus Rejection averaged across segmentation levels; a Segmentation main effect contrast with weights [1, −1, 1, −1], representing Short minus Long averaged across artifact handling levels; and a Segmentation × Artifact Handling interaction contrast with weights [1, −1, −1, 1]. Each contrast score was then transformed back toward the original SME scale using a signed square-root transformation, defined as the sign of the contrast value multiplied by the square root of its absolute value, which preserves the direction of the effect while returning values to a scale analogous to the original SME units. For each transformed contrast score distribution, normality was assessed using the Shapiro-Wilk test. Where normality was met, a one-sample t-test against zero was used; where normality was violated, a one-sample Wilcoxon signed-rank test against zero was used instead, with rank-biserial correlation reported as the effect size measure. Confidence intervals for the rank-biserial correlation were obtained using a percentile bootstrap, in which participants were resampled with replacement 10,000 times and the effect size was recomputed on each resample, with the 2.5th and 97.5th percentiles of the resulting distribution taken as the lower and upper bounds. Analyses were conducted separately for Cz and FCz. Holm correction was applied across the three contrast p-values within each electrode and outcome combination, as pairwise Spearman correlations between the three transformed contrast score distributions confirmed they were not independent.

Role of Trial Retention in bSME. To examine whether the bSME differences between pipelines were explained by trial retention differences, a contrast-to-contrast regression analysis was conducted using the participant-level contrast scores from the factorial decomposition and trial retention analyses. For each of the three planned orthogonal contrasts (Artifact Handling, Segmentation, and the interaction), the bSME contrast scores were regressed onto the corresponding trial count contrast scores using ordinary least squares regression, with each bSME outcome paired with its epoch-matched trial retention contrast scores.

For each regression, the OLS slope contribution was subtracted from each participant’s bSME contrast score, leaving adjusted scores that preserved the intercept of the regression. The intercept represents the predicted bSME contrast when the trial retention contrast is zero, that is, the baseline bSME difference that exists independently of trial retention. The adjusted scores were tested against zero using a Wilcoxon signed-rank test, with rank-biserial correlation reported as the effect size measure. Confidence intervals for the rank-biserial correlation were obtained using a percentile bootstrap. On each of 10,000 iterations, participants were resampled with replacement from the paired bSME and trial count contrast scores, the regression was refitted on the resampled data, and the adjusted scores and effect size were recomputed using that resample’s slope. The 2.5th and 97.5th percentiles of the resulting distribution were taken as the lower and upper bounds. Analyses were conducted separately for Cz and FCz. Holm correction was applied across the three contrast p-values within each electrode and outcome combination, as pairwise Spearman correlations between the three adjusted score distributions confirmed they were not independent.

## Results

### P50 Trial Retention

Table 2 shows the number of trials retained for P50 averaging across the four preprocessing pipelines, separately for S1 and S2. Descriptive statistics indicate clear differences in trial retention across both segmentation approach (Short vs Long) and artifact rejection strategy (Correction vs Rejection) for both stimulus epochs. Across S1 and S2, the Short-Correction pipeline retained the highest number of trials on average, followed closely by the Long-Correction pipeline, whereas both Rejection pipelines showed reduced trial retention, with the Long-Rejection pipeline retaining the fewest trials overall. One participant fell below the minimum trial threshold (< 40 trials) for the Long-Rejection pipeline.

**Table 1.** Four Preprocessing Pipelines Defined by Segmentation Strategy and Artifact Handling Method.

|  | **Correction** | **Rejection** |
| --- | --- | --- |
| **Short** | Single segmentation (-100 to 300ms); artifact rejection at midline electrodes followed by ocular correction using Gratton & Coles | Single segmentation (-100 to 300ms); threshold-based artifact rejection using $\pm 75\mu\text{V}$ criterion applied to all EEG and EOG channels |
| **Long** | Dual segmentation (-100 to 1000ms relative to S1); artifact rejection at midline electrodes followed by ocular correction using Gratton & Coles | Dual segmentation (-100 to 1000ms relative to S1); threshold-based artifact rejection using $\pm 75\mu\text{V}$ criterion applied to all EEG and EOG channels |

<sup>3</sup>The average reference was used instead of the mastoids due to a small post-auricular muscle response (PAMR) observed in some participants. Bell et al., (2004) recommend against using the mastoid reference when recording middle-latency ERPs due to potential interference from the PAMR. The average reference has been used in previous paired-click studies (Croft et al., 2004; Holstein et al., 2013; Rentzsch et al., 2008; Shen et al., 2020).
<sup>4</sup>Dual segments (S1 & S2 together) are baseline corrected to S1 prestimulus period prior to artifact rejection and S2 segments are re-baseline corrected to S2 prestimulus period prior to averaging.
<sup>5</sup>Artifact rejection procedure for the standard pipeline entailed the following: 1) Gradient check, maximal voltage steps of 10uV per millisecond; 2) $\pm 75\mu\text{V}$ amplitude threshold; 3) Low activity check, lowest allowed activity of 0.5uV across a 100ms interval.

**Table 2.** Trial Retention Per Preprocessing Pipeline.

|  | P50 trial counts |  |  |  |  |  |  |  |
| --- | --- | --- | --- | --- | --- | --- | --- | --- |
|  | Short Correction |  | Short Rejection |  | Long Correction |  | Long Rejection |  |
|  | No. trials |  | No. trials |  | No. trials |  | No. trials |  |
|  | S1 | S2 | S1 | S2 | S1 | S2 | S1 | S2 |
| **Average** | 120 | 119.98 | 111.71 | 111.57 | 119.98 | 119.98 | 101.07 | 101.07 |
| **SD** | 0 | 0.13 | 9.29 | 10.99 | 0.13 | 0.13 | 22.25 | 22.25 |
| **Max** | 120 | 120 | 120 | 120 | 120 | 120 | 120 | 120 |
| **Min** | 120 | 119 | 81 | 71 | 119 | 119 | 31 | 31 |
| **Removed** | 0 | 0 | 0 | 0 | 0 | 0 | 1 | 1 |

*Note.* Trials retained for S1 and S2 averages across each preprocessing pipeline. Values include the average, standard deviation (SD), minimum and maximum trials per pipeline, and number of participants removed due to insufficient retained trials (< 40 retained trials).

**Table 3.**
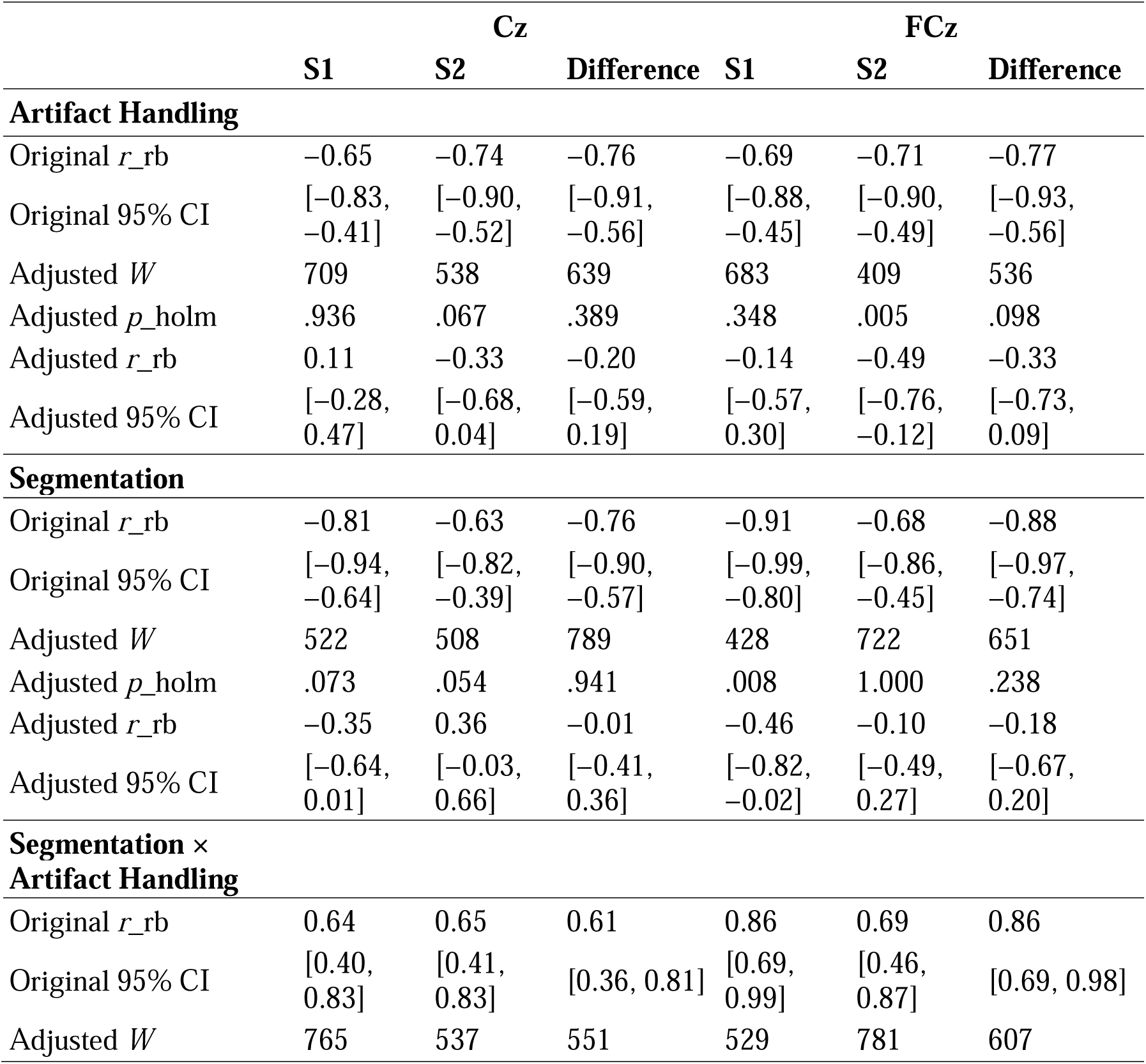
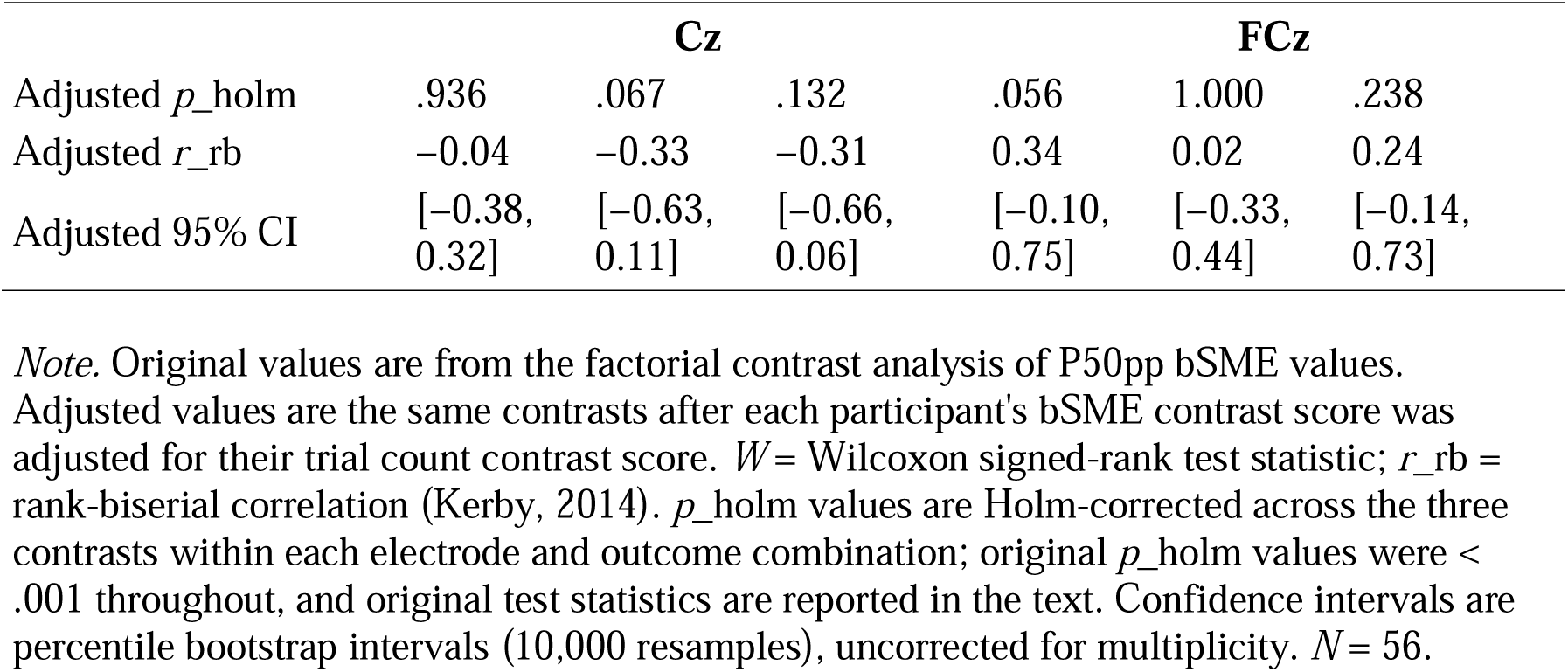
Original and Trial Retention-Adjusted bSME Contrast Effects at Cz and FCz.

The normality assumption was violated for all three contrast score distributions across both epochs (all *p* <.001), and Wilcoxon signed-rank tests were therefore used for all inferential tests. For S1, the Artifact Handling main effect was significant (*W* = 0, *p* <.001, *r*_rb = 1.00, 95% CI [1.00, 1.00]), indicating that Correction pipelines retained more trials than Rejection pipelines. The Segmentation main effect was also significant (*W* = 0, *p* <.001, *r*_rb = 1.00, 95% CI [1.00, 1.00]), indicating that Short segmentation retained more trials than Long segmentation. The Segmentation × Artifact Handling interaction was significant (*W* = 0, *p* <.001, *r*_rb = −1.00, 95% CI [−1.00, −1.00]), reflecting that the Correction advantage over Rejection was larger under Long segmentation than under Short segmentation. Results for S2 were consistent with S1: the Artifact Handling main effect (*W* = 0, *p* <.001, *r*_rb = 1.00, 95% CI [1.00, 1.00]), Segmentation main effect (*W* = 3, *p* <.001, *r*_rb = 1.00, 95% CI [0.98, 1.00]), and Segmentation × Artifact Handling interaction (*W* = 3, *p* <.001, *r*_rb = −1.00, 95% CI [−1.00, −0.98]) were all significant, with the same directional pattern observed.

### Standardized Measurement Error (SME)

#### RMS-bSME: Group-Level Data Quality

Across S1, S2, and the difference score, the Short-Correction (S1: 0.31 µV, SE = 0.02; S2: 0.30 µV, SE = 0.02; Difference: 0.44 µV, SE = 0.02) and Long-Correction (S1: 0.32 µV, SE = 0.02; S2: 0.30 µV, SE = 0.02; Difference: 0.44 µV, SE = 0.02) pipelines showed the lowest RMS-bSME values and were nearly indistinguishable from one another. The Short-Rejection pipeline produced slightly higher values (S1: 0.33 µV, SE = 0.02; S2: 0.31 µV, SE = 0.02; Difference: 0.45 µV, SE = 0.02), while the Long-Rejection pipeline consistently yielded the highest RMS-bSME values across all three outcomes (S1: 0.36 µV, SE = 0.02; S2: 0.35 µV, SE = 0.02; Difference: 0.50 µV, SE = 0.03). Ratio score RMS-bSME values were substantially inflated across all pipelines (raw ratio range: 8.60–8.90 µV; see Appendix C Table 4 and Table 5 for full values) due to division by small S1 amplitudes and were therefore not further analysed.

**Table 4.** Ratio Score RMS-bSME Values Per Preprocessing Pipeline at Cz.

| Pipeline | Ratio<br>RMS-bSME | Ratio<br>mean | Ratio<br>SD | Log<br>ratio<br>RMS-bSME | Log<br>ratio<br>mean | Log<br>ratio<br>SD | Capped<br>ratio<br>RMS-bSME | Capped<br>ratio<br>mean | Capped<br>ratio<br>SD |
| --- | --- | --- | --- | --- | --- | --- | --- | --- | --- |
| **Short-Correction** | 8.60 | 4.96 | 7.03 | 1.17 | 2.05 | 0.51 | 0.44 | 0.41 | 0.17 |
| **Long-Correction** | 8.70 | 5.02 | 7.10 | 1.16 | 1.05 | 0.51 | 0.44 | 0.41 | 0.17 |
| **Short-Rejection** | 8.83 | 4.81 | 7.41 | 1.17 | 1.06 | 0.49 | 0.44 | 0.41 | 0.16 |
| **Long-Rejection** | 8.90 | 5.05 | 7.32 | 1.19 | 1.08 | 0.50 | 0.45 | 0.42 | 0.16 |

*Note.* RMS-bSME, mean, and SD values for raw ratio, log-transformed ratio, and capped ratio P50pp bSME outcomes across all four preprocessing pipelines at Cz. Raw ratio values are substantially inflated due to division by small S1 P50 amplitudes.

**Table 5.**
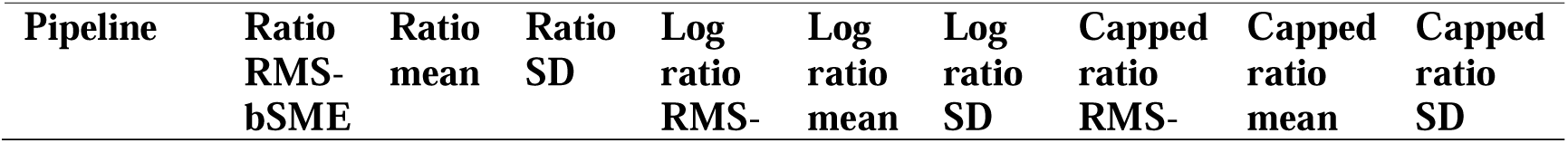
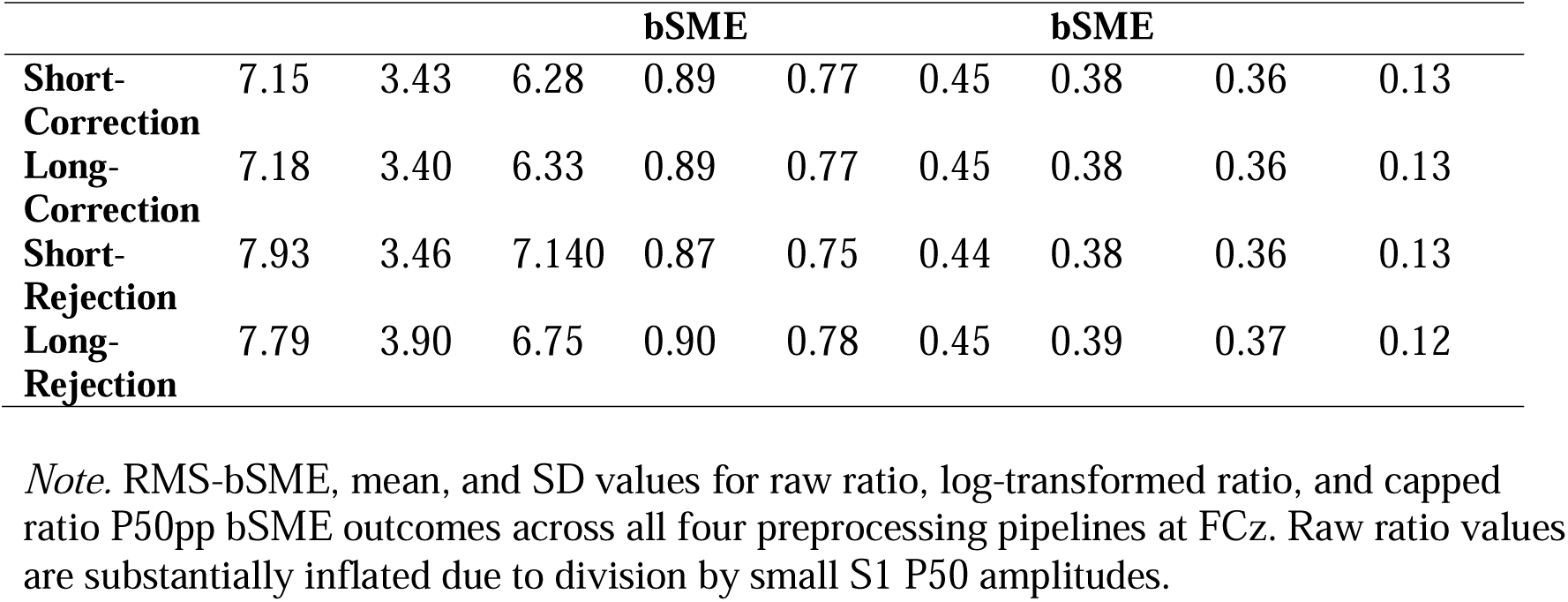
Ratio Score RMS-bSME Values Per Preprocessing Pipeline at FCz.

RMS-bSME values at FCz followed the same pattern across all three outcomes (see Appendix A, Figure 5). The Short-Correction (S1: 0.37 µV, *SE* = 0.02; S2: 0.35 µV, *SE* = 0.02; Difference: 0.51 µV, *SE* = 0.02) and Long-Correction (S1: 0.37 µV, *SE* = 0.02; S2: 0.35 µV, *SE* = 0.02; Difference: 0.51 µV, *SE* = 0.02) pipelines again showed the lowest RMS-bSME values, followed by Short-Rejection (S1: 0.39 µV, *SE* = 0.02; S2: 0.36 µV, *SE* = 0.01; Difference: 0.53 µV, *SE* = 0.02), with the Long-Rejection pipeline yielding the highest values overall (S1: 0.44 µV, *SE* = 0.03; S2: 0.39 µV, *SE* = 0.02; Difference: 0.59 µV, *SE* = 0.03).

#### Factorial Decomposition of Pipeline Differences

All contrast score distributions violated the normality assumption at Cz (all Shapiro-Wilk *p* <.05), and Wilcoxon signed-rank tests were therefore used for all three contrasts across all outcomes. For S1, the Artifact Handling main effect was significant (*W* = 282, *p* <.001, *r*_rb = −0.65, CI [−0.83, −0.41]), indicating that Correction pipelines produced lower bSME values than Rejection pipelines on the variance scale. The Segmentation main effect was also significant (*W* = 151, *p* <.001, *r*_rb = −0.81, CI [−0.94, −0.64]), indicating that Short segmentation produced lower bSME values than Long segmentation. The Segmentation × Artifact Handling interaction was significant (*W* = 287, *p* <.001, *r*_rb = 0.64, CI [0.40, 0.83]), reflecting that the Rejection disadvantage was disproportionately larger under Long segmentation than under Short segmentation. Results for S2 were consistent with S1: the Artifact Handling main effect (*W* = 210, *p* <.001, *r*_rb = −0.74, CI [−0.90, −0.52]), Segmentation main effect (*W* = 297, *p* <.001, *r*_rb = −0.63, CI [−0.82, −0.39]), and Segmentation × Artifact Handling interaction (*W* = 282, *p* <.001, *r*_rb = 0.65, CI [0.41, 0.83]) were all significant, with the same directional pattern observed. For the difference score, all three contrasts were again significant: Artifact Handling (*W* = 191, *p* <.001, *r*_rb = −0.76, CI [−0.91, −0.56]), Segmentation (*W* = 189, *p* <.001, *r*_rb = −0.76, CI [−0.90, −0.57]), and the interaction (*W* = 313, *p* <.001, *r*_rb = 0.61, CI [0.36, 0.81]). The Cz results are displayed in figure 6 below.

Results at FCz mirrored those at Cz across all three outcomes (see figure 7). All contrast score distributions violated normality (all Shapiro-Wilk *p* <.05), and Wilcoxon signed-rank tests were used throughout. For S1, the Artifact Handling main effect (*W* = 250, *p* <.001, *r*_rb = −0.69, CI [−0.88, −0.45]), Segmentation main effect (*W* = 70, *p* <.001, *r*_rb = −0.91, CI [−0.99, −0.80]), and interaction (*W* = 109, *p* <.001, *r*_rb = 0.86, CI [0.69, 0.99]) were all significant, with the same directional pattern as Cz. For S2, the Artifact Handling main effect (*W* = 228, *p* <.001, *r*_rb = −0.71, CI [−0.90, −0.49]), Segmentation main effect (*W* = 259, *p* <.001, *r*_rb = −0.68, CI [−0.86, −0.45]), and interaction (*W* = 250, *p* <.001, *r*_rb = 0.69, CI [0.46, 0.87]) were all significant. For the difference score, all three contrasts were significant: Artifact Handling (*W* = 187, *p* <.001, *r*_rb = −0.77, CI [−0.93, −0.56]), Segmentation (*W* = 99, *p* <.001, *r*_rb = −0.88, CI [−0.97, −0.74]), and interaction (*W* = 113, *p* <.001, *r*_rb = 0.86, CI [0.69, 0.98]).

#### Role of Trial Retention in bSME

The results of the adjusted score tests are displayed in Table 3 in Appendix D. The table shows the results of the original bSME factorial contrast analysis alongside the results obtained after trial retention differences across pipelines were accounted for.

Artifact Handling contrast. At Cz, the artifact handling contrast was not significant for S1, S2, or the difference score. At FCz, the artifact handling contrast was significant for S2, in the same direction as the original finding, with a reduced effect size relative to the original, but not for S1 or the difference score. Across both electrodes, the artifact handling effect on bSME was not consistent across outcomes after trial retention was accounted for, with a significant result in the same direction as the original finding observed only for S2 at FCz.

Segmentation contrast. At Cz, the segmentation contrast was not significant for S1, S2, or the difference score. At FCz, the segmentation contrast was significant for S1, with Short segmentation still producing lower bSME values than Long segmentation, consistent with the original direction, with a reduced effect size relative to the original, but not for S2 or the difference score. After trial retention was accounted for, the segmentation effect was significant in the same direction as the original finding only for S1 at FCz.

Interaction contrast. At Cz, the interaction contrast was not significant for S1, S2, or the difference score. At FCz, the interaction contrast was not significant for S1, S2, or the difference score. After trial retention was accounted for, none of the interaction contrasts were significant at either electrode across any outcome.

Overall pattern. Every one of the 18 contrasts was reduced in magnitude relative to the original factorial contrast analysis, and 16 were no longer significant after trial retention was accounted for. The two that remained significant, the artifact handling effect for S2 at FCz (*r*_rb = −0.49, 95% CI [−0.76, −0.12]) and the segmentation effect for S1 at FCz (*r*_rb = −0.46, 95% CI [−0.82, −0.02]), were both in the same direction as the original findings but substantially reduced in magnitude (original *r*_rb = −0.71 and −0.91, respectively). This pattern indicates that trial retention is the primary contributor to the bSME differences observed between pipelines, with the number of trials retained for averaging accounting for the large majority of the pipeline differences in measurement error across outcomes and electrodes.

## Discussion

Taken together, the trial retention and bSME analyses provide a coherent account of how preprocessing pipelines can influence data quality in the P50 paired-click paradigm. Across both analyses, a consistent pattern emerged in which Correction pipelines outperformed Rejection pipelines. Both analyses revealed a significant Segmentation × Artifact Handling interaction, reflecting that the two factors do not operate independently, with the Rejection disadvantage being larger under Long segmentation than under Short segmentation.

### The Role of Trial Retention

Trial retention analyses showed that Correction pipelines retained significantly more trials than Rejection pipelines across both S1 and S2. SME analyses converged on the same pattern across all three analytical approaches. RMS-bSME values were lowest for the Short-Correction and Long-Correction pipelines, which were nearly indistinguishable from one another, and highest for the Long-Rejection pipeline, with Short-Rejection producing intermediate values. Consistent with the approach from Luck et al., (2021) to connect the SME to the effect size and statistical power, the bSME values were squared before computing the contrast scores in the factorial contrast analysis. The factorial contrast findings corroborated the RMS-bSME findings, with significant Artifact Handling and Segmentation main effects and a significant interaction at both Cz and FCz across all outcomes; Short and Long Correction pipelines produced the highest precision of P50 estimates.

After trial retention was accounted for in the regression analysis, 16 of the 18 bSME pipeline contrasts were non-significant, indicating that trial retention is the primary contributor to the observed bSME differences between pipelines. The two surviving effects, the artifact handling effect for S2 at FCz and the segmentation effect for S1 at FCz, were both in the same direction as the original findings with reduced effect sizes, further supporting the conclusion that the pipeline differences in bSME are largely driven by differences in trial retention. Therefore, this study recommends that future paired-click studies utilize EEG preprocessing pipelines that maximize trial retention through artifact correction methods in order to maximize the precision of P50 estimates and derived gating metrics.

### Artifact Handling: Correct or Reject Ocular Artifacts?

Our findings provide empirical support for both Correction pipelines as the most favourable preprocessing options, as they consistently produced the highest trial retention and the highest precision across all outcomes at both electrodes. As stated above, this outcome was predominantly driven by trial retention differences. As noted by previous researchers, there are several problems with threshold-based artifact rejection; it does not remove small VEOG artifacts that can affect results (Croft & Barry, 2000), it can result in a substantial loss of EEG data (Krishnaveni et al., 2005), and studies have shown that this method is inferior to correction methods (O’Toole & Iacono, 1987; Verleger et al., 1982). Researchers have therefore recommended against the use of threshold-based artifact rejection to handle ocular-related artifacts (Croft & Barry, 2000). We support such recommendations, showing that rejecting trials contaminated by artifacts leads to a significant reduction in trial retention and P50 data quality (increased measurement error). Thus, in line with the calls for standardization of paired-click methodology (de Wilde et al., 2007; Patterson et al., 2008), this study recommends that future paired-click studies should utilize a combined approach of non-ocular artifact rejection and ocular correction to maximize trial retention and P50 data quality.

However, it should be noted that there is a known limitation of regression-based ocular correction methods. The regression-based ocular correction method used in this study, Gratton and Coles, can result in a slight change in ERP amplitudes. This is caused by forward propagation, where neural activity in the EEG channels propagates forward to the EOG channels that are used in the regression model. Croft and Barry (2002) distinguish a calculation phase, in which the correction coefficient is estimated, from a subtraction phase, in which the weighted EOG is removed from the EEG. Assuming the calculation phase produces an accurate coefficient, they showed that forward propagation can introduce an error in the subtraction phase because the EOG activity that is subtracted from the EEG signal contains a portion of genuine neural activity. Subtracting the EOG activity therefore removes some real EEG along with the ocular artifact. Previous paired-click researchers have posited that since the P50 component is small (∼1–3 µV), this error-driven amplitude change could distort the P50 ratio scores (Hall et al., 2006). However, it has been shown that this subtraction-phase error results in minimal amplitude changes of auditory N100 and P200 components at central sites where the P50 is also measured from (Croft & Barry, 2002). Note that with a realistic regression coefficient of.1, the actual amount of EEG removed from the EEG-electrode signal is only 1% of the total.

Although the error is minimal, there are other ocular correction methods that are available (see Ronca et al., 2024 for a tutorial review). One of these correction methods, independent component analysis (ICA), has been previously examined within the SME framework. In a similar approach to this study, Zhang et al., (2024a) found that combining ICA ocular correction with the rejection of non-ocular extreme artifacts improved SME for several ERP components, especially amplitude scores. This study chose a regression-based method due to the availability of external EOG template channels, which enables a simple and effective approach to correcting ocular artifacts (Ronca et al., 2024). However, future studies should empirically examine and compare the different ocular correction methods that are available to determine which method produces the most precise estimates of P50 measures.

### Segmentation strategy: Short vs Long

While Short and Long Correction pipelines produced similar trial retention and bSME values for P50 analysis, the regression analysis provides modest evidence that Short segmentation produces lower bSME values than Long segmentation independently of trial retention. Of the 6 segmentation contrasts, only the FCz S1 contrast remained significant after trial retention was accounted for, with a substantially reduced effect size relative to the original finding. This suggests that trial retention accounts for the large majority of the segmentation effect, but the surviving result nonetheless points towards a small residual data quality advantage for Short segmentation that cannot be fully explained by trial retention differences alone.

While the regression analysis provides modest evidence of a residual data quality advantage for Short segmentation, there is no strong empirical evidence that Long segmentation provides any data quality benefit over Short segmentation. As stated previously, Long segmentation maintains true matched pairs by ensuring that any trial containing a non-ocular artifact in either S1 or S2 is rejected entirely. Short segmentation rejects S1 and S2 segments independently, which can result in unmatched averages. Given that trial-to-trial variability in P50 responses has been shown to influence averaged P50 metrics and derived measures (Patterson et al., 2000), maintaining matched pairs may be theoretically advantageous. However, since trial retention is the primary driver of bSME values (precision), as demonstrated by the regression analysis, and until studies empirically compare the effects of matched versus unmatched P50 averages on the S1-S2 difference score and gating ratio, the pipeline that retains the most trials remains the more defensible recommendation. Given calls for standardisation of methodological practices in the paired-click paradigm literature (de Wilde et al., 2007; Patterson et al., 2008), this study recommends future paired-click studies utilize the Short segmentation approach.

## Filter settings

This study has noted the importance of two other preprocessing choices in P50 data analysis, filter settings and choice of reference. As demonstrated in Figure 1, the P50 component is abolished under 0.5-30Hz filter settings, as the N100 activity is allowed into the signal and significantly drags down the P50 waveform. This supports recommendations from Chang et al. (2012), who demonstrated that filter settings directly influence P50 amplitude, signal-to-noise ratio, and measurement stability. A 10-50Hz bandpass filter is therefore recommended for P50 analysis, as the 10Hz high-pass cutoff is necessary to attenuate the lower-frequency N100 activity that would otherwise obscure the P50 component.

### Reference electrode

Reference electrode choice represents a further methodological consideration in the paired-click paradigm. Although the mastoid reference is commonly used (Dalecki et al., 2015, 2016; Kisley et al., 2004; Lijffijt et al., 2009; Oranje et al., 2006; Rosburg et al., 2009; Yadon et al., 2009), this study used the average reference due to a postauricular muscle response (PAMR) that was observed in the mastoid channels (see Appendix B, figure 8). As shown in Figure 8, when using the mastoid reference, even a small PAMR amplitude (∼1µV) was sufficient to induce an artifactual peak at approximately 16ms. This early component preceded the P30 component and displayed a posterior topographical distribution that is inconsistent with a genuine auditory component. Using the average reference, no artifactual peak preceding the P30 is observed.

One acknowledged drawback of the average reference is that midline components such as the P50 will be smaller in amplitude compared to a mastoid reference. However, the average reference has been used in previous paired-click studies (Croft et al., 2004; Holstein et al., 2013; Rentzsch et al., 2008; Shen et al., 2020; Thoma et al., 2020). Furthermore, the PAMR is triggered by loud sounds and the optimal tone intensity to elicit stable P50 responses is 85-90 dB (de Wilde et al., 2007), making it likely a small PAMR response will be observed in some participants. Therefore, this study supports previous recommendations to avoid the mastoid reference when recording middle-latency auditory ERPs (Bell et al., 2004). In keeping with the broader aim of advancing methodological standardisation in the paired-click paradigm (de Wilde et al., 2007; Patterson et al., 2008), we recommend that future studies use the average reference where viable, particularly when a high-density electrode setup is available, and to provide explicit justification where an alternative reference is used.

## Conclusion

In conclusion, the present study provides empirical support for specific preprocessing recommendations for the P50 paired-click paradigm. Regarding artifact handling, the findings support the use of ocular correction methods, combined with rejection of non-ocular artifacts at midline electrodes, over the commonly used threshold-based rejection procedures applied to all EEG and EOG channels. As noted above, threshold-based rejection of ocular artifacts has previously been cautioned against (Croft & Barry, 2000). Our findings show correction-based pipelines consistently produced higher trial retention and lower measurement error than rejection-based pipelines across all outcomes at both electrodes. Further recommendations are proposed in relation to segmentation strategy, filter setting and choice of reference. Together, these recommendations align with previous calls for standardisation of preprocessing practices in the paired-click literature (de Wilde et al., 2007; Patterson et al., 2008), and the present findings provide an empirical basis for moving towards more consistent and transparent reporting of preprocessing decisions in P50 paired-click studies.

## Appendix

### Appendix A

**Figure 5.**
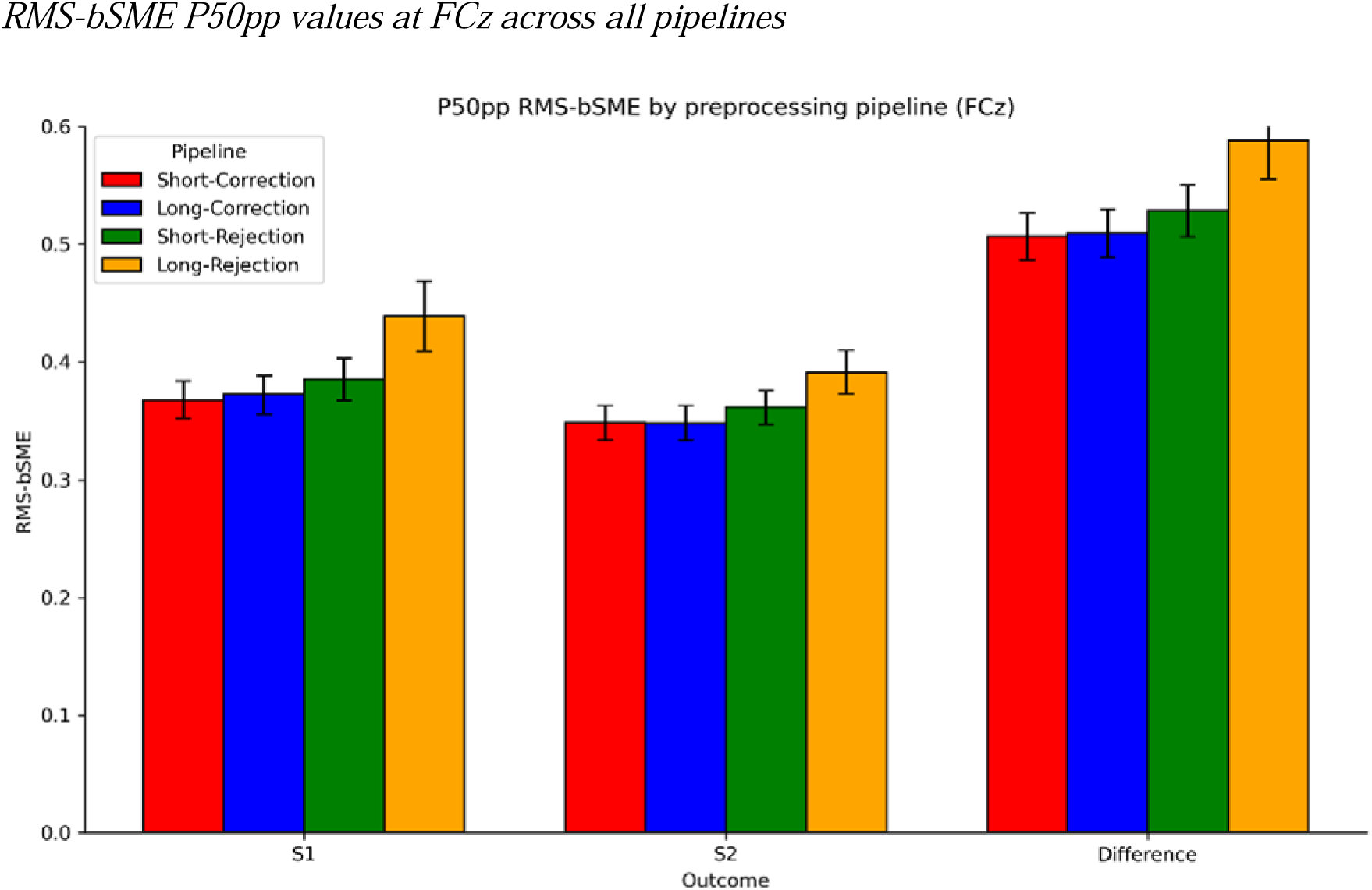
RMS-bSME P50pp values at FCz across all pipelines. *Note.* All pipelines are shown: Short Correction (red), Long Correction (blue), Short Rejection (green), and Long Rejection (orange). Error bars represent the standard error of the RMS-bSME for each pipeline.

**Figure 6.**
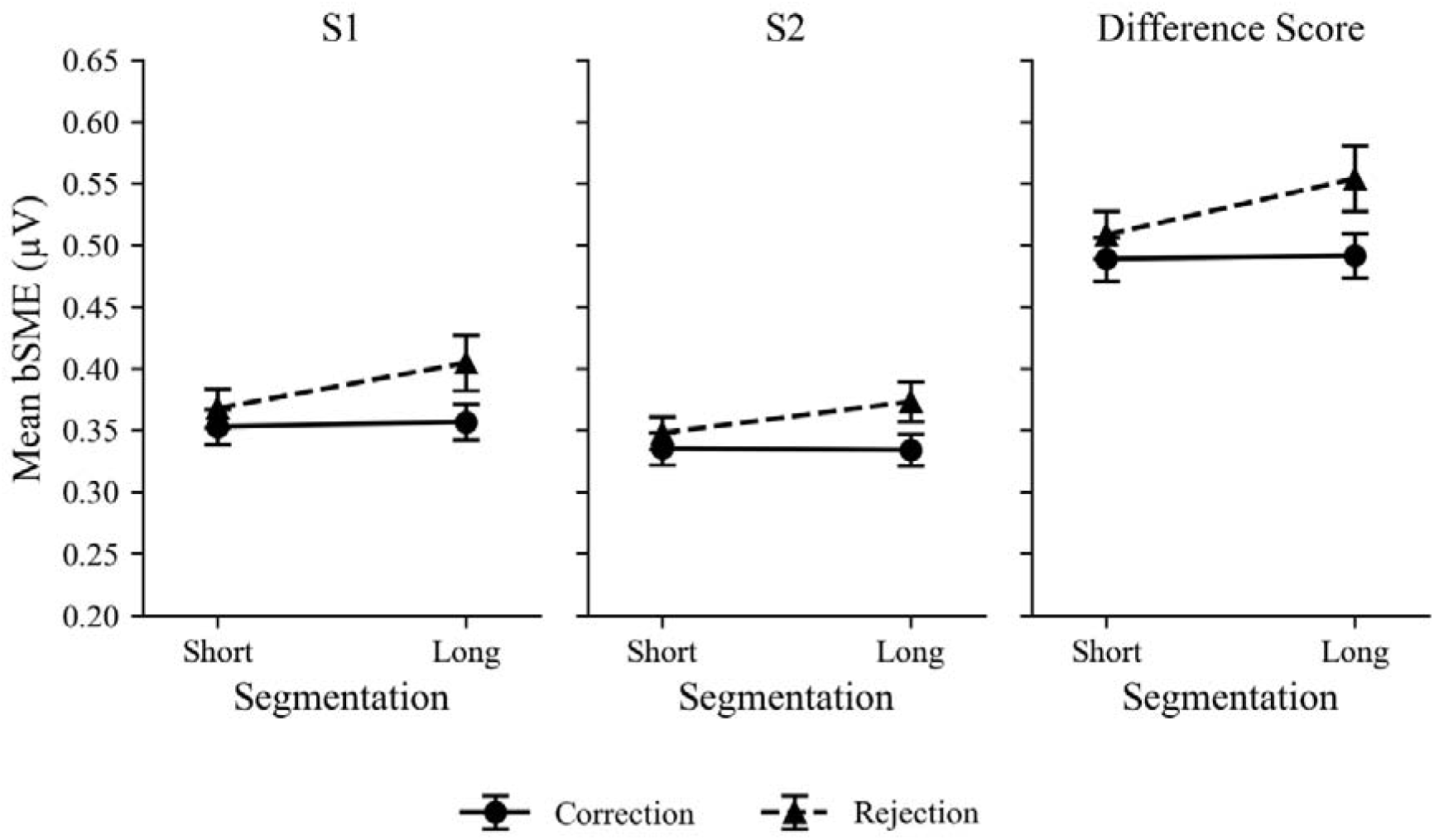
Mean P50pp bSME Values at Cz Across Preprocessing Pipelines. *Note.* Mean bootstrap SME values for S1, S2, and the difference score at Cz across Segmentation levels (Short vs Long) separately for Correction and Rejection artifact handling strategies. Error bars represent the standard error of the mean bSME value within each pipeline.

**Figure 7.**
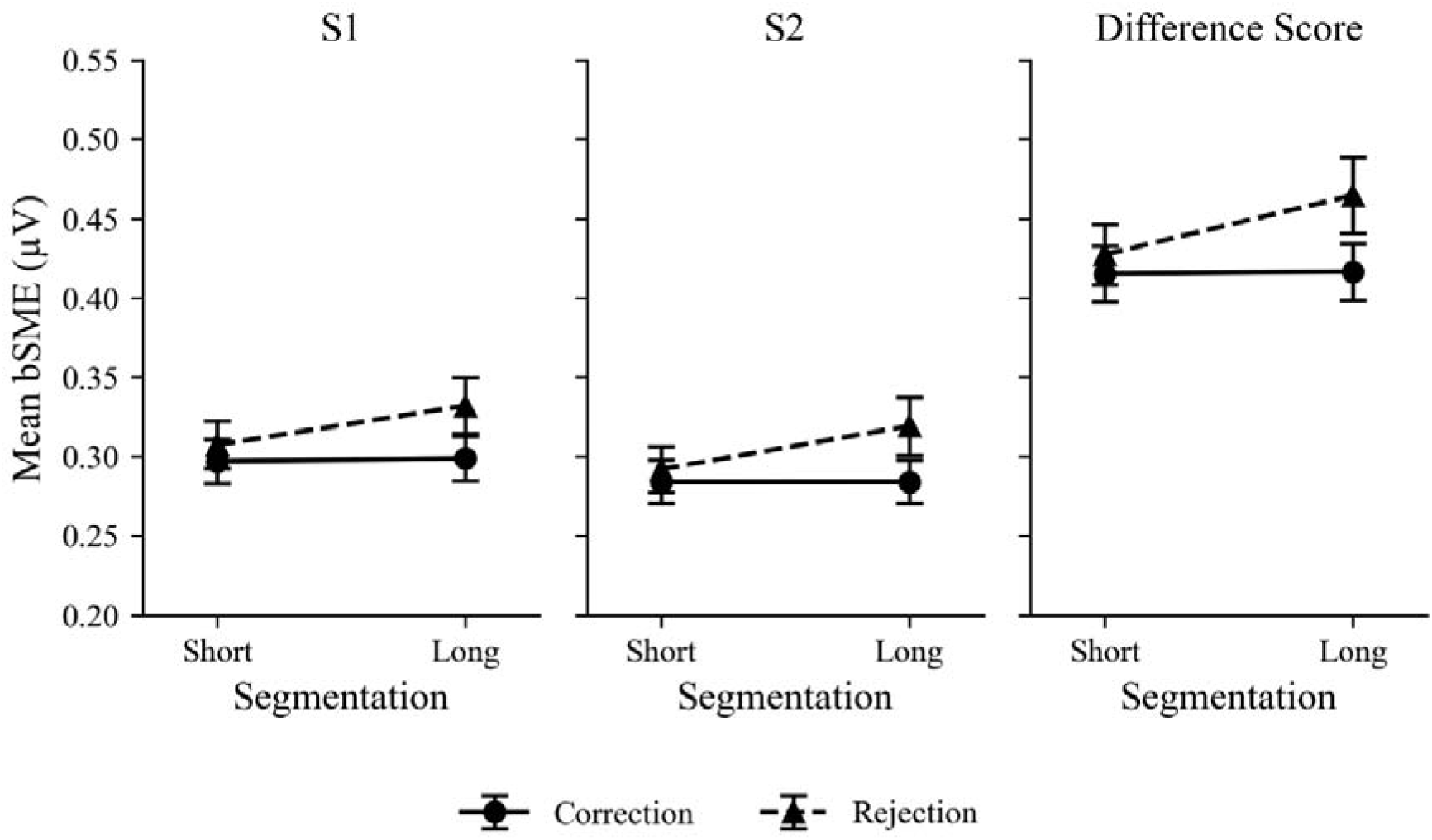
Mean P50pp bSME Values at FCz Across Preprocessing Pipelines. *Note.* Mean bootstrap SME values for S1, S2, and the difference score at FCz across Segmentation levels (Short vs Long) separately for Correction and Rejection artifact handling strategies. Error bars represent the standard error of the mean bSME value within each pipeline.

### Appendix B

**Figure 8.**
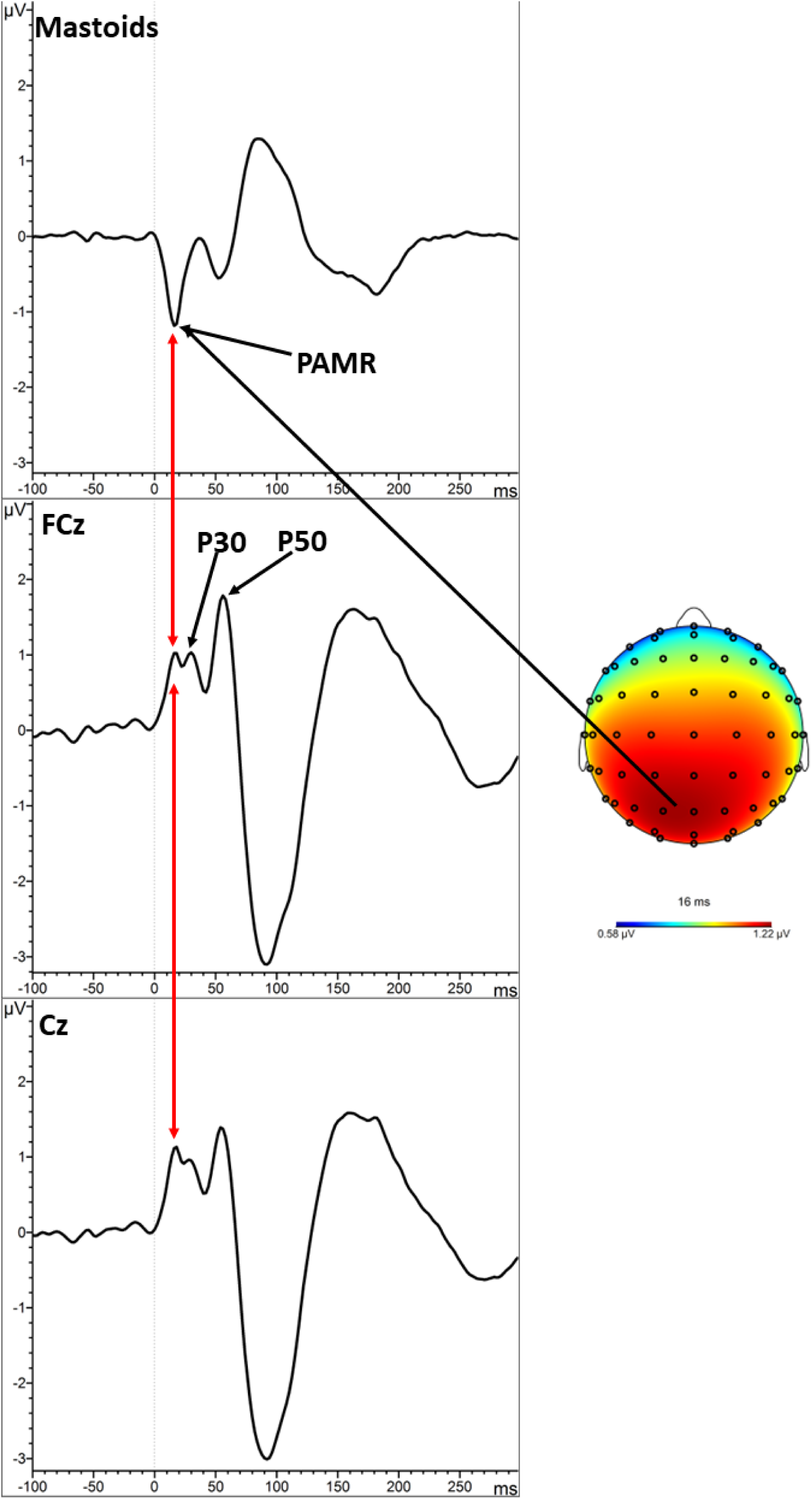
Grand Average Localizer (S1 & S2) showing the PAMR at the Mastoids, Cz, and FCz. *Note.* Grand average ERP waveforms at the Mastoids (top), FCz (middle), and Cz (bottom) for the Short-Correction pipeline using the mastoid reference, averaged across S1 and S2 trials. The red arrow indicates the post-auricular muscle response (PAMR) peak at approximately 16ms, which is present at the mastoid electrodes and is projected onto FCz and Cz when the mastoids are used as the reference electrode, appearing immediately prior to the P30 component. The topographical distribution map at 16ms shows a posterior maximum, which is inconsistent with the frontocentral distribution expected for a genuine auditory component. P30 and P50 components are labelled on the FCz waveform for reference.

### Appendix C

#### Ratio scores

Ratio score bSME values were substantially inflated across all pipelines and electrodes relative to the S1, S2, and difference score outcomes, with raw ratio RMS-bSME values ranging from 7.15 to 8.90 compared to a range of 0.30 to 0.59 for the other outcomes (see Appendix C, Table 4 and Table 5 for full values). This inflation arises from division by small S1 P50 amplitudes, which produces unstable and disproportionately large ratio values at the individual participant level. Log-transformed and capped ratio (scores above 2 capped to 2) variants were less severely inflated but remained substantially higher than the non-ratio outcomes. Ratio scores are therefore not recommended as a primary outcome measure for bSME analyses in the paired-click paradigm.

### Appendix D

#### Appendix E: Sound Calibration Procedure

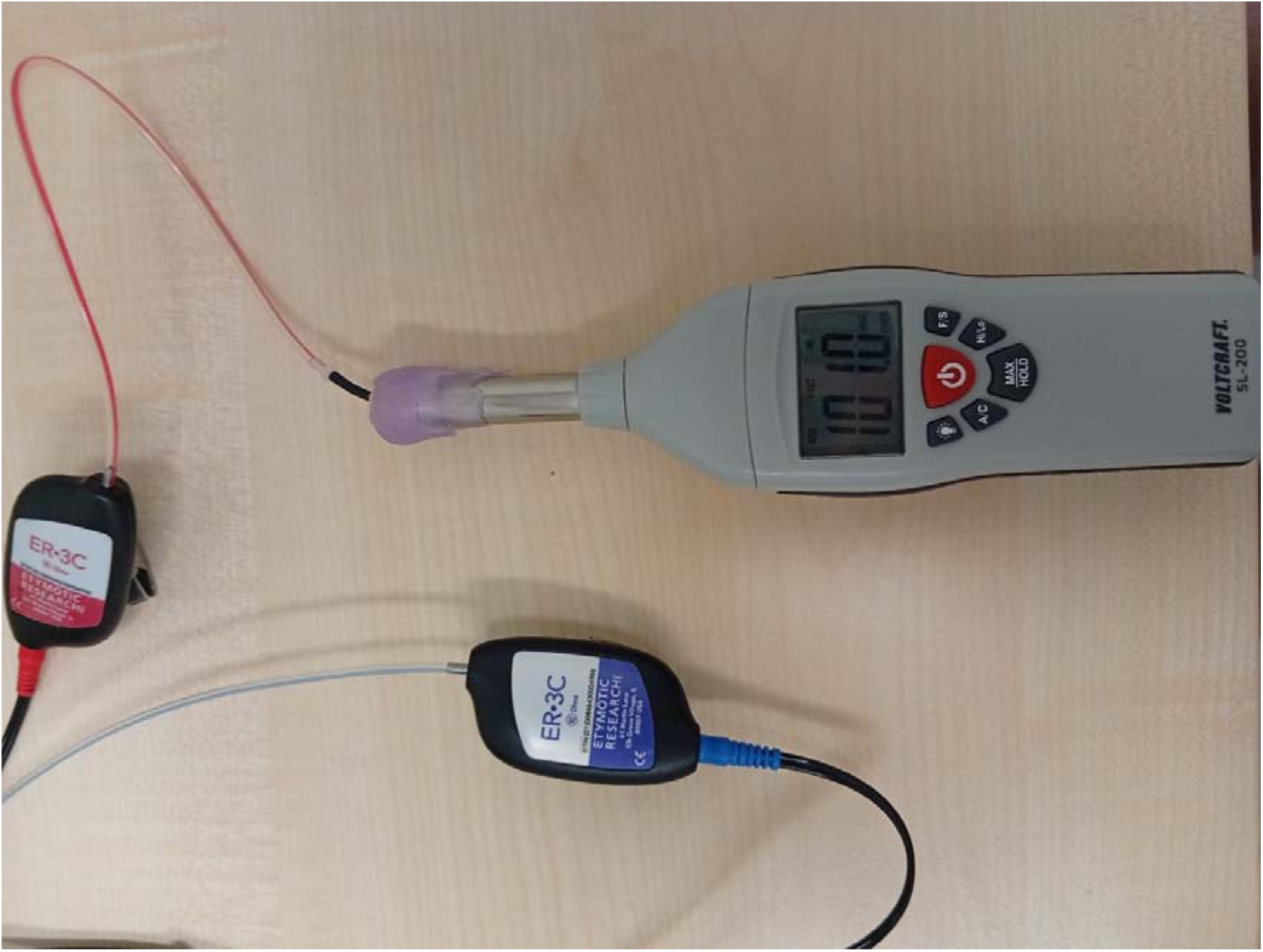

To calibrate the 1000 Hz auditory tones to be 100 dB SPL, an airtight seal was used to connect the earphone (ER-3C insert earphones, Etymotic) to the sound meter (Voltcraft SL-200). An airtight seal was used to replicate the airtight seal that is created when the foam eartip is inserted into the ear of the participant. The sound meter settings were set to dBA, Fast, High, Max. 500ms 1000 Hz tones were used and volume was modulated until 100 dBA was achieved. To note, when the airtight seal was removed and the eartip was placed against the sound meter, the tone intensity was 80 dBA.

## Footnotes

1 Studies that have used single segmentation approach: Bak et al., (2011); Boutros et al., (1999, 2004, 2009); Chang et al., (2012); Clementz et al., (1998); Croft et al., (2004); Dalecki et al., (2011); Davies et al., (2009); Fuerst et al., (2007); Gjini et al., (2011); Hall et al., (2006); Kisley et al., (2004); Knott et al., (2009); Lijffijt et al., (2009); Shen et al., (2020); Yadon et al., (2009).

2 Studies that have used dual segmentation approach: Chien et al., (2019); Dalecki et al., (2015, 2016); Grunwald et al., (2003); Holstein et al., (2013); Micoulaud-Franchi et al., (2015); Rentzsch et al., (2008); Thoma et al., (2020); Thomas et al., (2010); White & Yee, (2006).

3 The average reference was used instead of the mastoids due to a small post-auricular muscle response (PAMR) observed in some participants. Bell et al., (2004) recommend against using the mastoid reference when recording middle-latency ERPs due to potential interference from the PAMR. The average reference has been used in previous paired-click studies (Croft et al., 2004; Holstein et al., 2013; Rentzsch et al., 2008; Shen et al., 2020)

4 Dual segments (S1 & S2 together) are baseline corrected to S1 prestimulus period prior to artifact rejection and S2 segments are re-baseline corrected to S2 prestimulus period prior to averaging

5 Artifact rejection procedure for the standard pipeline entailed the following: 1) Gradient check, maximal voltage steps of 10uV per millisecond; 2) ±75uV amplitude threshold; 3) Low activity check, lowest allowed activity of 0.5uV across a 100ms interval.

